# Natural variation in the control of flowering and shoot architecture in diploid *Fragaria* species

**DOI:** 10.1101/2021.12.22.473817

**Authors:** Guangxun Fan, Javier Andrés, Klaus Olbricht, Elli Koskela, Timo Hytönen

## Abstract

In perennial fruit and berry crops of the Rosaceae family, flower initiation occurs in late summer or autumn after downregulation of a strong repressor *TERMINAL FLOWER1* (*TFL1*) and flowering and fruiting takes place the following growing season. Rosaceous fruit trees typically form two types of axillary shoots, short flower-bearing shoots called spurs and long shoots that are respectively analogous to branch crowns and stolons in strawberry. However, regulation of flowering and shoot architecture differs between species and environmental and endogenous controlling mechanisms have just started to emerge. In woodland strawberry (*Fragaria vesca* L.), long days maintain vegetative meristems and promote stolon formation by activating *TFL1* and *GIBBERELLIN 20-OXIDASE4* (*GA20ox4*), respectively, while silencing of these factors by short days and cool temperatures induces flowering and branch crown formation. We characterized flowering responses of 14 accessions of seven diploid *Fragaria* species native to diverse habitats in the northern hemisphere, and selected two species with contrasting environmental responses, *F. bucharica* Losinsk. and *F. nilgerrensis* Schlecht. ex J. Gay for detailed studies together with *F. vesca*. Similar to *F. vesca, F. bucharica* was induced to flower in short days at 18°C and regardless of photoperiod at 11°C after silencing of *TFL1. F. nilgerrensis* maintained higher *TFL1* expression level and likely required cooler temperatures or longer exposure to inductive treatments to flower. We also found that high expression of *GA20ox4* was associated with stolon formation in all three species, and its downregulation by short days and cool temperature caused branch crown formation in *F. vesca* and *F. nilgerrensis*, although the latter did not flower. *F. bucharica*, in contrast, rarely formed branch crowns, regardless of flowering or *GA20ox4* expression level. Our findings highlighted diploid *Fragaria* species as a rich source of genetic variation controlling flowering and plant architecture, with potential applications in breeding of Rosaceous crops.

## 1.1 Introduction

The Rosaceae family contains economically important perennial crops, ranging from herbaceous species such as strawberries (*Fragaria* spp.), to fruit trees like apples (*Malus × domestica* Borkh.) or peaches (*Prunus persica* (L.) Batsch) (Kurokura et al., 2013). In strawberries and Rosaceous fruit trees, floral induction takes place during summer or autumn, and flower initials continue developing until late autumn. As the season advances towards winter, these species gradually enter a period of dormancy that is broken after a genetically determined period of cold temperatures called chilling requirement. After that, when the growing season begins in the spring, vegetative growth resumes and blooming occurs (Wilkie et al., 2008; Bangerth, 2009; Kurokura et al., 2013; Costes et al., 2014). Similarities in the seasonal growth cycles of strawberries and Rosaceous fruit trees suggest that species such as woodland strawberry (*F. vesca* L.) and other diploid strawberries may be successfully used as models for studying developmental events during the seasonal cycle.

The cues required for floral induction and subsequent floral initiation differ from species to species, and may depend on environmental, developmental, cultural or genetic factors. For instance, in apple and sweet cherry (*Prunus avium* L.), floral initiation depends on temperature, with species-specific optima (Sønsteby and Heide, 2019; Heide et al., 2020). In the perennial herbaceous model species *F. vesca*, floral induction is highly dependent on the interaction of temperature and photoperiod in autumn, and natural populations exhibit differences in their responses to these environmental cues. In some populations, cool temperature of 9°C is sufficient to induce flowering independently of photoperiod, whereas in other populations grown at cool temperature, the promotive effect of short days (SD) is still evident (Heide and Sønsteby, 2007). Moreover, a population sampled from the North of Norway shows a strikingly altered yearly growth cycle with an obligatory requirement for vernalization (Heide and Sønsteby, 2007; Koskela et al., 2017). Given that such variation in environmental responses exists within a single strawberry species, it is imaginable that extending these studies to other related diploid strawberry species could reveal further adaptations to the local environment. Characterizing the available diversity within the *Fragaria* genus may prove useful not only for researchers but also for breeders looking for novel breeding targets to improve climatic adaptation of Rosaceous crops.

As developmental events depend on the timing of meristem differentiation and the meristematic fate itself, studies on meristem fate may provide insights into climatic adaptation of plants. In Rosaceous species, shoot apical meristems (SAMs) located at shoot tips can either generate new vegetative tissues, develop terminal inflorescences, or in some species abort spontaneously (Costes et al., 2014). Meristems located in leaf axils develop axillary buds (AXBs), which can remain latent, develop into vigorously growing long shoots or into short shoots with limited extension growth. Short shoots (also known as spurs or dwarf shoots) are characterized by short internode length and a rosette-like appearance. In many Rosaceous species, terminal meristems borne on short shoots are more prone to receiving the floral induction stimulus and initiating flowers than meristems borne on vigorously growing long shoots (Wilkie et al., 2008; Sønsteby and Heide, 2019). Therefore, the balance between short and long shoots defines the yield potential and affects the choice and expenses of cultural practices such as pruning or training.

In the herbaceous perennial model species *F. vesca* and the cultivated strawberry *F.* × *ananassa* (Duch.), AXBs can remain latent, develop into branch crowns (BCs) analogous to short shoots, or into stolons that resemble the vigorously growing long shoots in Rosaceous trees (Hytönen et al., 2004; Heide and Sønsteby, 2007). As strawberry inflorescences are borne terminally at the apices of the primary crown and BCs, the number or BCs has a direct effect on yield potential. On the other hand, stolons are indispensable for clonal reproduction of strawberry cultivars. Being such important processes for determining the feasibility of strawberry growing, genetic regulation of floral initiation and AXB fate have received the attention of several researchers during the last decade.

Molecular studies in both *F. vesca* (Koskela et al., 2012; Rantanen et al., 2015) and *F.* × *ananassa* (Koskela et al., 2016) have highlighted the role of *TERMINAL FLOWER1* (*TFL1*) as a floral repressor. In *F. vesca*, the *CONSTANS* (*FvCO*) - *FLOWERING LOCUS T1* (*FvFT1*) - *SUPPRESSOR OF OVEREXPRESSION OF CONSTANS1* (*FvSOC1*) photoperiodic pathway activates *FvTFL1* in long days (LDs), and flower induction occurs in SDs after gradual downregulation of *FvTFL1* (Koskela et al., 2012; Mouhu et al., 2013; Rantanen et al., 2015; Kurokura et al., 2017). However, this photoperiodic pathway regulates flowering only within a narrow temperature range between 13 to 20°C (Rantanen et al., 2015). Lower temperatures repress *FvTFL1* and induce flowering independently of photoperiod, while higher temperatures activate *FvTFL1* and inhibit flowering regardless of the photoperiod (Rantanen et al., 2015).

The function of *TFL1* in Rosaceous species outside the *Fragaria* genus is very conserved. Silencing or knocking out *TFL1* homologs in apple and pear (*Pyrus communis* L.) result in reduced juvenility, precocious flowering and even perpetual flowering (Kotoda et al., 2006; Flachowsky et al., 2012; Freiman et al., 2012; Charrier et al., 2019). Likewise, loss-of-function of *TFL1* homologs in roses (*Rosa* spp.) and *F. vesca* also lead to perpetual flowering (Iwata et al., 2012; Koskela et al., 2012; Bai et al., 2021). The seasonal expression pattern of *TFL1* is highly connected to the yearly growth cycle in apple, *F. vesca* and roses, as *TFL1* is activated in the SAM during the vegetative growth phase and downregulated before the floral induction to allow flower initiation (Mimida et al., 2011; Iwata et al., 2012; Kurokura et al., 2013; Koskela et al., 2017). With such a conserved function and expression patterns across Rosaceous species, it is reasonable to expect that results from studies on *TFL1* in one species are applicable to other species.

Regulation of AXB fate in Rosaceae has been mainly studied at the phenotype level, perhaps because the molecular processes taking place within the well-protected AXB are difficult to examine. However, a recent reports in *F. vesca* demonstrated that stolon development in this species requires *GIBBERELLIN 20-OXIDASE4* (*FvGA20ox4*) that is activated within the AXBs under LD conditions via an *FvSOC1*-dependent photoperiodic pathway at 18°C (Mouhu et al., 2013; Tenreira et al., 2017; Andrés et al., 2021). Higher temperature of 22°C upregulates *FvGA20ox4* independently of *FvSOC1*, whereas at cooler temperature (11°C), *FvGA20ox4* is de-activated in both SD and LD conditions (Andrés et al., 2021). Cool temperatures, as well as SDs at 18°C, promote BC development instead of stolons in the seasonal flowering *F. vesca*, and although the same environmental cues induce flowering, these two processes can occur independently (Andrés et al., 2021). However, the fate of the youngest AXB located immediately below the SAM is directly dependent on the vegetative/generative status of the SAM; if the SAM is induced to flower, the youngest AXB develops a branch crown to continue the growth of the plant in a sympodial fashion (Sugiyama et al., 2004; Andrés et al., 2021). Based on similar phenotypic effects from growth regulator applications (Ramina et al., 1985; Reekie and Hicklenton 2002; Webster, 2004; Hytönen et al., 2009), similar mechanisms may be involved in the control of AXB development to short or long shoots in different Rosaceous species.

Studying natural variation present in related species is based on the rationale that species sampled from diverse environments have faced different selection pressures, leading to evolution of local adaptation and phenotypical differences. Recently, local adaptation was studied in wild species and landrace accessions of peach from a wide geographical range in China, and candidate genes related to flowering time and adaptation to climate change were identified (Li et al., 2021).

We adopted the approach to study environmental responses in wild diploid species of *Fragaria* originating from a wide geographical range. Our collection of wild diploid *Fragaria* included accessions native to high altitude (*F. bucharica* and *F. nubicola* Lindl. from the Himalayas) as well as accessions endemic to less harsh environments (*F. iinumae* Makino growing in the Japanese archipelago and *F. nilgerrensis* and *F. pentaphylla* Losinsk. from South East Asia). In addition, we included our Finnish reference accession of *F. vesca* that is adapted to temperate climate, *F. viridis* Weston from Central Europe and *F. chinensis* Losinsk. from North Western China. We analyzed the collection in terms of flowering habits and vegetative development under controlled environment and selected *F. bucharica* and *F. nilgerrensis* for detailed analysis together with *F. vesca*. Our results indicated that *TFL1* homologs are key integrators of temperature and photoperiodic cues in these three species and that altered regulation of *TFL1* may explain variation in their flowering habits. We also found that the activities of *GA20ox4* homologs correlated with AXB fate in the studied diploid strawberry species.

## 1.2 Materials and Methods

### 1.2.1 Plant Material

Seasonal flowering accession FIN56 (PI551792, National Clonal Germplasm Repository, Corvallis, OR, USA) of *F. vesca* was used as a control. Other 14 accessions of seven wild diploid strawberries were provided by the Professor Staudt Collection, Germany (Supplementary Table 1). All plant materials used in this study were propagated from stolon cuttings in the greenhouse. Plants were first grown on jiffy pellets (Jiffy Products International) for three weeks, and then transplanted to 8 × 8 cm pots with fertilized peat (Kekkilä, Finland). Liquid fertilizer (Kekkilä, N-P-K: 17-4-25, Finland) was given to the plants biweekly.

### 1.2.2 Treatments and observations

Plants were grown in a greenhouse under 18h LD at 18°C for four weeks before the experiments started. In greenhouse, plants were illuminated by natural light, and high-pressure sodium lamps (Airam 400W, Kerava, Finland) at a photosynthetic photon flux density (PPFD) of 120 μmol m^−2^ s^−1^ were used to extend the day length. Temperature and photoperiod treatments (11°C and 18°C) were carried out in growth chambers equipped with LED lamps (AP67, Valoya, Finland; 200 μmol m^−2^ s^−1^ of PPFD). The treatment details of each experiment are described in the figure legends. During the experiment, the number of leaves, stolons and BCs were observed weekly, and stolons were removed after recording. For flowering time observations, both the numbers of leaves from the primary leaf rosette and the number of days when the first fully open flower emerged were recorded. In this study, the BC number only referred to the number of axillary leaf rosettes, and the sympodial BC arising from the topmost axil upon floral initiation was not counted.

### 1.2.3 Gene expression data

Shoot apex samples were collected for gene expression analysis. Total RNA was extracted as described by Koskela et al., (2012) and treated with rDNase (Macherey-Nagel GmbH, Düren, Germany) according to the manufacturer’s instructions. cDNA was synthesized from 500 ng of total RNA using ProtoScript II Reverse Transcriptase according to manufacturer’s instructions (New England Biolabs). SYBR Green I master mix was used for quantitative real-time PCR (qRT-PCR) in a total reaction volume of 10 μland analyzed by LightCycler 480 instrument (Roche) as described by Koskela et al., (2012). Four biological replicates and three technical replicates were used for qRT-PCR analysis using the primers listed in the Supplementary Table 2. Relative expression levels were calculated by ΔΔCt method as described by Pfaffl, (2007). *FvMSI1* (*MULTICOPY SUPPRESSOR OF IRA1*) was used as a reference gene for normalization.

### 1.2.4 Statistical analyses

Either logistic regression or ANOVA was conducted to test the main factors, and pairwise comparisons were performed by TukeyHSD. The statistical analyses were done using R.4.1.0 (R Core Team, 2021), the stats (v4.1.0; R Core Team, 2021) and the DescTools (v0.99.42; Andri et mult. al. S, 2021) packages.

## 1.3 Results

### 1.3.1 Flowering time variation among diploid strawberry species

To get the first insights into flowering responses to environmental cues in different diploid strawberry species, 14 accessions of seven species were subjected to 18h LD and 12h SD treatments at 11°C for six weeks, followed by flowering observations in LDs at 18°C. Eleven accessions showed clear flowering response to at least one of the treatments (Figure 1A). All the plants of two *F. bucharica* and three *F. viridis* accessions flowered after both LD and SD treatments, showing a similar photoperiod-independent flowering response to cool temperature as SD genotypes of *F. vesca* (Heide and Sønsteby 2007; Rantanen et al. 2015). In the two *F. chinensis* accessions, floral induction took place in all the LD-grown plants, whereas only roughly half of the plants were induced under SD conditions. Opposite photoperiodic response was found in two accessions of *F. nilgerrensis*, in which SDs resulted in a higher percentage of flowering plants. In *F. nilgerrensis* #1, as well as in *F. iinumae* #1 and *F. nubicola* #1, flowering occurred only after the SD treatment (Figure 1A). Finally, flowering was not observed in *F. nubicola* #2 and the two *F. pentaphylla* accessions under either photoperiod.

**Figure 1.**
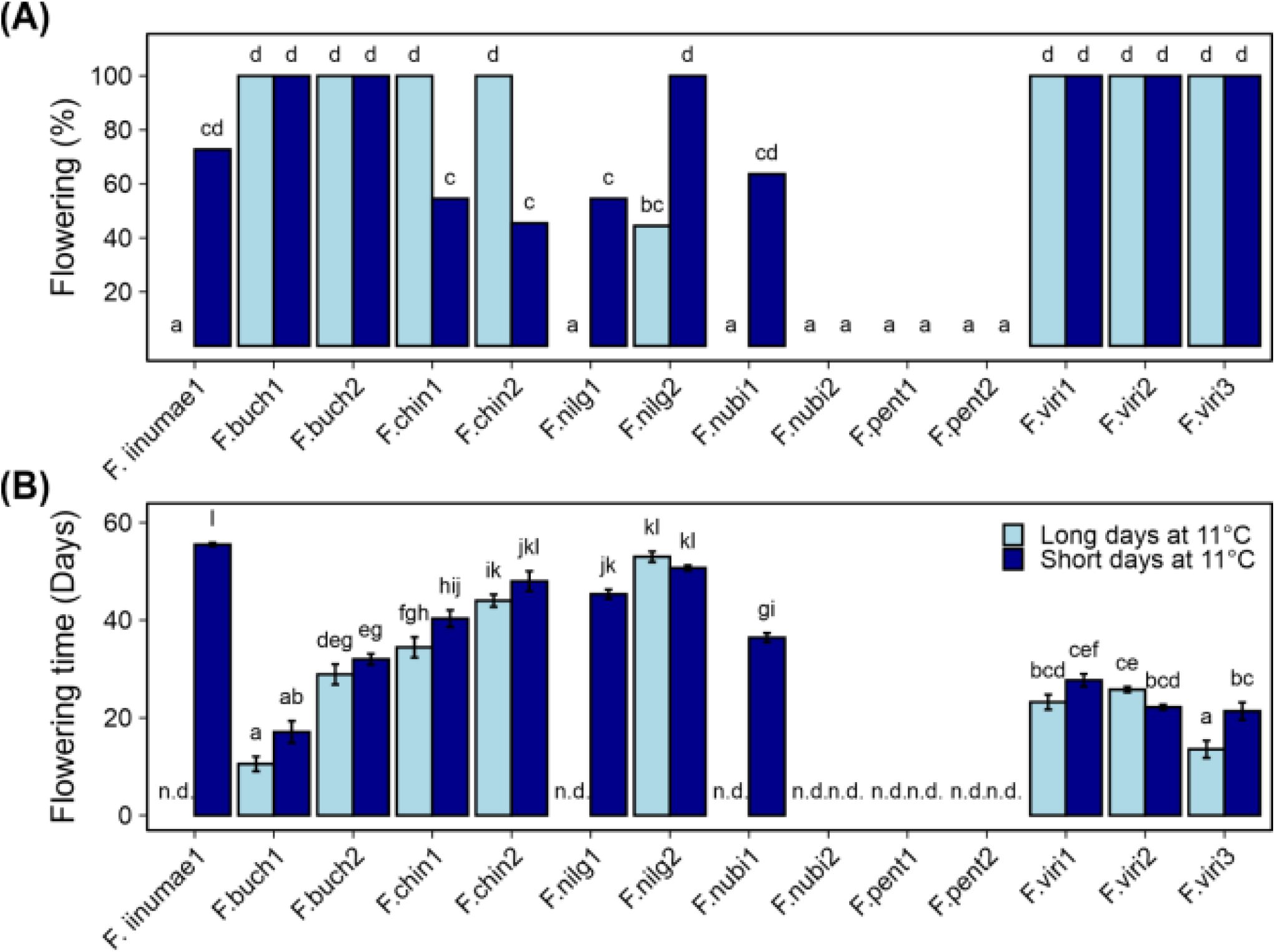
Flowering responses of diploid species to 12h short day (SD) and 18h long day (LD) treatments at 11°C. Percentage of flowering plants **(A)** and days to the first open flower **(B)**. Stolon-propagated plants were grown under 18 or 12-h photoperiod at 11°C for 6 weeks followed by 10 weeks under 18h photoperiod at 18°C. Flowering was recorded every other day, starting after the treatments. Error bars represent the standard error of the mean (n=10) and different letters indicate significant differences calculated by ANOVA and Tukey’s test (P < 0.05). “n.d.” = flowering not observed during the course of the experiment (16 weeks).

There were also significant differences in flowering time between the species and accessions (Figure 1B). Accessions of *F. bucharica* and *F. viridis* flowered rapidly after the temperature treatment independently of photoperiod, except for *F. viridis* #3 in which LDs promoted flowering. On the other hand, *F. iinumae* and *F. nilgerrensis* accessions flowered significantly later than those two fast flowering species.

Because we were not able to induce *F. nubicola* #2 and the two *F. pentaphylla* accessions to flower under SDs or LDs at 11°C, we tested if a prolonged cold treatment could induce them to flower. After one month in 12h SD at 14-15°C, plants were moved to 5-6°C for about 4 months. This treatment induced flowering in all the plants of both *F. pentaphylla* accessions that did not flower after 11°C treatment (Supplementary Table 3). However, *F. nubicola* #2 did not flower, and inductive conditions for this accession remained an enigma. Furthermore, *F. iinumae* #1 did not flower, although it flowered in a previous experiment.

### 1.3.2 The effect of photoperiod on flowering time and AXB fate at 18°C

The screening experiment with 14 *Fragaria* accessions revealed an interesting diversity of flowering responses. Therefore, we were curious to find out how the species would behave in direct comparison with the seasonal flowering *F. vesca* reference genotype, FIN56. We selected *F. bucharica* #1 because it flowered first in the initial screening experiment and *F. nilgerrensis* #1 because it flowered late and had a clear photoperiodic response at 11°C (Figure 1; the rest of the experiments included only one genotype per accession and therefore *F. bucharica* #1 and *F. nilgerrensis* #1 are referred to as *F. bucharica* and *F. nilgerrensis* from here onwards). First, we decided to subject the three species to LDs or SDs at 18°C, because the phenotypical responses and gene expression profiles in *F. vesca* are well characterized under these conditions.

*F. vesca* plants were induced to flower by a six-week period of SDs at 18°C, while the LD-grown control plants did not flower. *F. bucharica* had a similar photoperiodic response, although the SD-grown *F. bucharica* flowered significantly earlier than SD-grown *F. vesca*, both in terms of flowering time and leaf number at flowering. In contrast, *F. nilgerrensis* did not flower under either photoperiod at 18°C (Figure 2).

**Figure 2.**
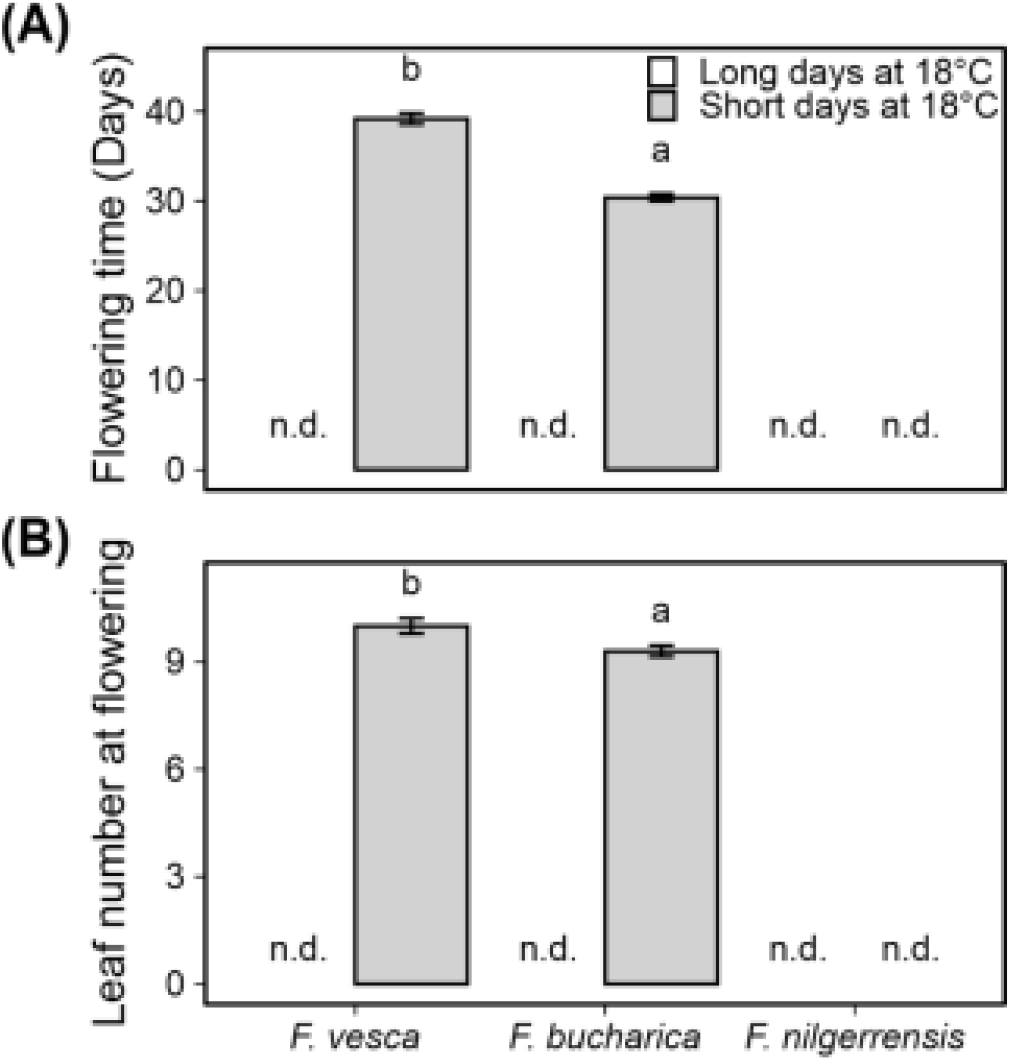
Flowering time in three diploid strawberry species grown in SDs or LDs at 18°C. Number of days (A) and number of leaves developed (B) until the first flower opened. Stolon-propagated plants were grown in LDs (18-h) or SDs (12-h) at 18°C for six weeks, followed by ten more weeks in LDs (18-h) at 18°C. Flowering time was recorded every other day and leaf number was scored at flowering time. Error bars represent the standard error of the mean (n=10), and different letters indicate significant differences calculated by ANOVA and Tukey’s test (P < 0.05). “n.d.” = flowering not observed during the course of the experiment (16 weeks).

At 18°C, *F. vesca* ceased stolon development after three weeks of SDs. The final number of stolons in *F. vesca* at the end of the experiment was significantly higher in LDs than in SDs (Figure 3A). *F. bucharica* did not stop stolon development under either photoperiod, although LDs slightly promoted stolon development also in this species (Figure 3A). Stolon development in *F. nilgerrensis* ceased after five weeks in SDs at 18°C, whereas LDs promoted stolon development until the end of the experiment (Figure 3A).

**Figure 3.**
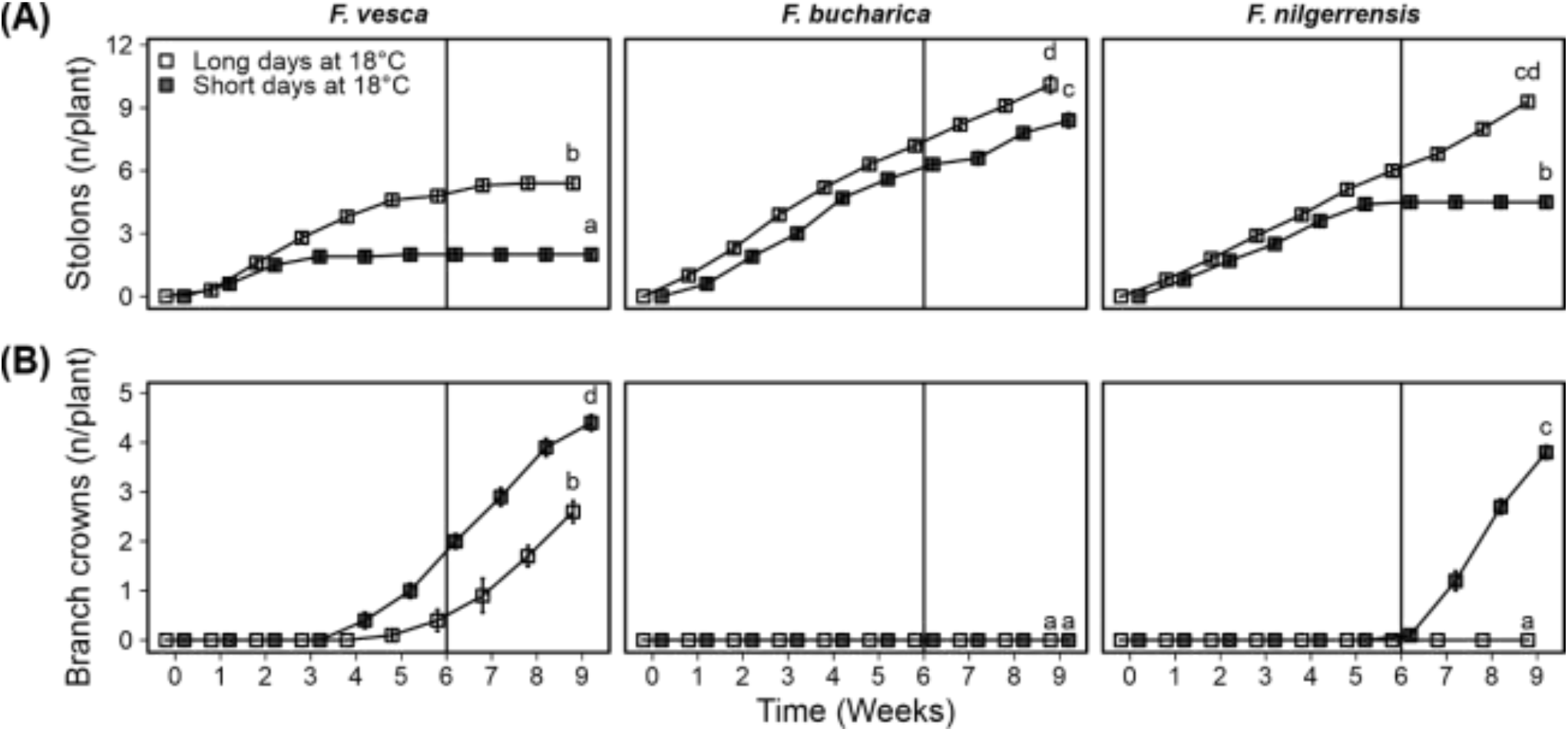
Axillary bud fate of three diploid strawberry species grown in SDs or LDs at 18°C. Number of stolons **(A)** and branch crowns **(B)** per plant. Stolon-propagated plants were grown in LDs (18-h) or SDs (12-h) at 18°C for six weeks, followed by three more weeks in LDs (18-h) at 18°C. Number of stolons and branch crowns was recorded weekly. Error bars represent the standard error of the mean (n=10) and different letters indicate significant differences calculated by ANOVA and Tukey’s test (P < 0.05).

The three species behaved very differently in terms of BC development at 18°C. In *F. vesca*, BC development was observed in both photoperiods, with SDs promoting BC development (Figure 3B). Intriguingly, *F. bucharica* did not develop any BCs under either photoperiod. In *F. nilgerrensis*, six weeks of SDs at 18°C strongly promoted BC development whereas the LD-grown *F. nilgerrensis* did not develop any BCs, indicating a clear photoperiodic effect (Figure 3B).

### 1.3.3 Altered expression of key genes is associated with different phenotypical responses at 18°C

To gain an initial idea of how the photoperiodic pathway functions in *F. bucharica* and *F. nilgerrensis*, we decided to study the expression of *SOC1, TFL1* and *GA20ox4* in these species in comparison to the reference accession of *F. vesca*. The *SOC1* genes had a very clear photoperiodic response in all three species and the expression patterns were very similar. *SOC1* remained active in LDs at 18°C, whereas SDs downregulated the gene, especially in *F. nilgerrensis* (Figure 4A). Our results suggest that the photoperiodic regulation of *SOC1* at 18°C is conserved between these diploid species.

**Figure 4.**
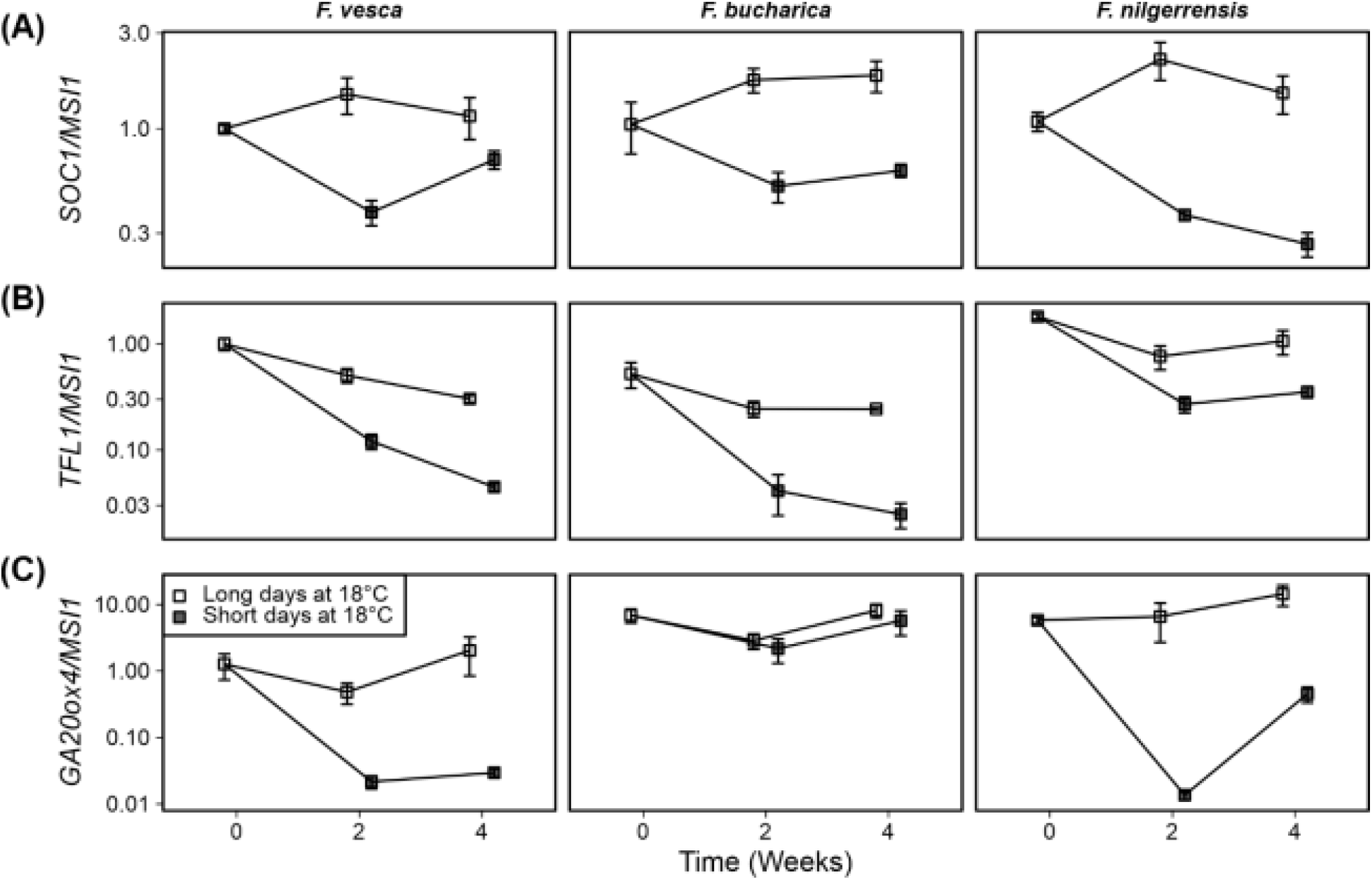
Gene expression patterns in three diploid strawberry species grown under SDs or LDs at 18°C. *SOC1* expression **(A)**; *TFL1* expression **(B)**; *GA20ox4* expression **(C)** in shoot apical samples. Stolon-propagated plants were grown in LDs (18-h) or SDs (12-h) at 18°C for six weeks. Shoot apical samples were collected at the beginning of the treatments, and 2 and 4 weeks later. Week 0 *F. vesca* samples were used as a calibrator for relative expression analysis. Error bars represent the standard error of the mean (n = 3–4).

Earlier experiments in *F. vesca* have shown that *FvTFL1* is gradually de-activated after transferring the plants to SD conditions (Koskela et al., 2012). In our current experiment, *TFL1* expression dropped to low levels in both *F. vesca* and *F. bucharica* within two weeks under SD conditions, and the expression further declined until week four (Figure 4B). In *F. nilgerrensis, TFL1* expression in SDs remained at higher level than in the other two species, although there was still a clear photoperiodic effect. Our expression data suggests that *TFL1* is expressed at constitutively higher level in *F. nilgerrensis* than in *F. vesca* or *F. bucharica* at 18°C, and that the high activity of *TFL1* may inhibit floral induction in the experimental conditions used. Moreover, regulation of *TFL1* in *F. nilgerrensis* (*FnTFL1*) does not follow the expression pattern of *SOC1*, implying that *FnTFL1* is regulated by factor(s) other than the *SOC1*-dependent pathway.

In *F. vesca, FvGA20ox4* was rapidly downregulated upon exposure to SDs and a similar expression pattern was observed in *F. nilgerrensis*, with clear downregulation of *GA20ox4* by week two under SDs, but not under LDs (Figure 4C). Concurring with the stolon phenotype, we found that the expression of *GA20ox4* in *F. bucharica* remained at a high level in LDs and SDs at 18°C. The finding in *F. bucharica* suggests that in this species the regulation of *GA20ox4* is uncoupled from the expression of *SOC1*, leading to continuous and photoperiod-independent development of stolons at 18°C.

### 1.3.4 Three diploid *Fragaria* species exhibit distinct phenotypic responses at 11°C

Next, we wanted to study the phenotypical responses of the three *Fragaria* species at 11°C at a more detailed level. We repeated the photoperiodic experiment at cooler temperature of 11°C and observed flowering and AXB fates by counting the number of stolons and branch crowns regularly. It has been earlier shown in *F. vesca* that cool temperature induces flowering independently of photoperiod (Heide and Sønsteby, 2007; Rantanen et al., 2015; Andrés et al., 2021). We found similar photoperiod-independent response in both *F. vesca* and *F. bucharica* after 6-week treatments at 11°C with significantly later flowering time in *F. bucharica*, while the control plants grown under LD conditions at 18°C remained vegetative until the end of the experiment (Figure 5). By contrast, no flowering plants were observed in *F. nilgerrensis* in this experiment, although 50% plants flowered in the initial screening under SDs at 11°C. In further experiments in *F. bucharica*, more than half of the plants were induced to flower already after 2-week treatments at 11°C, and four weeks was enough to induce flowering in almost all plants (Supplementary Figure 1).

**Figure 5.**
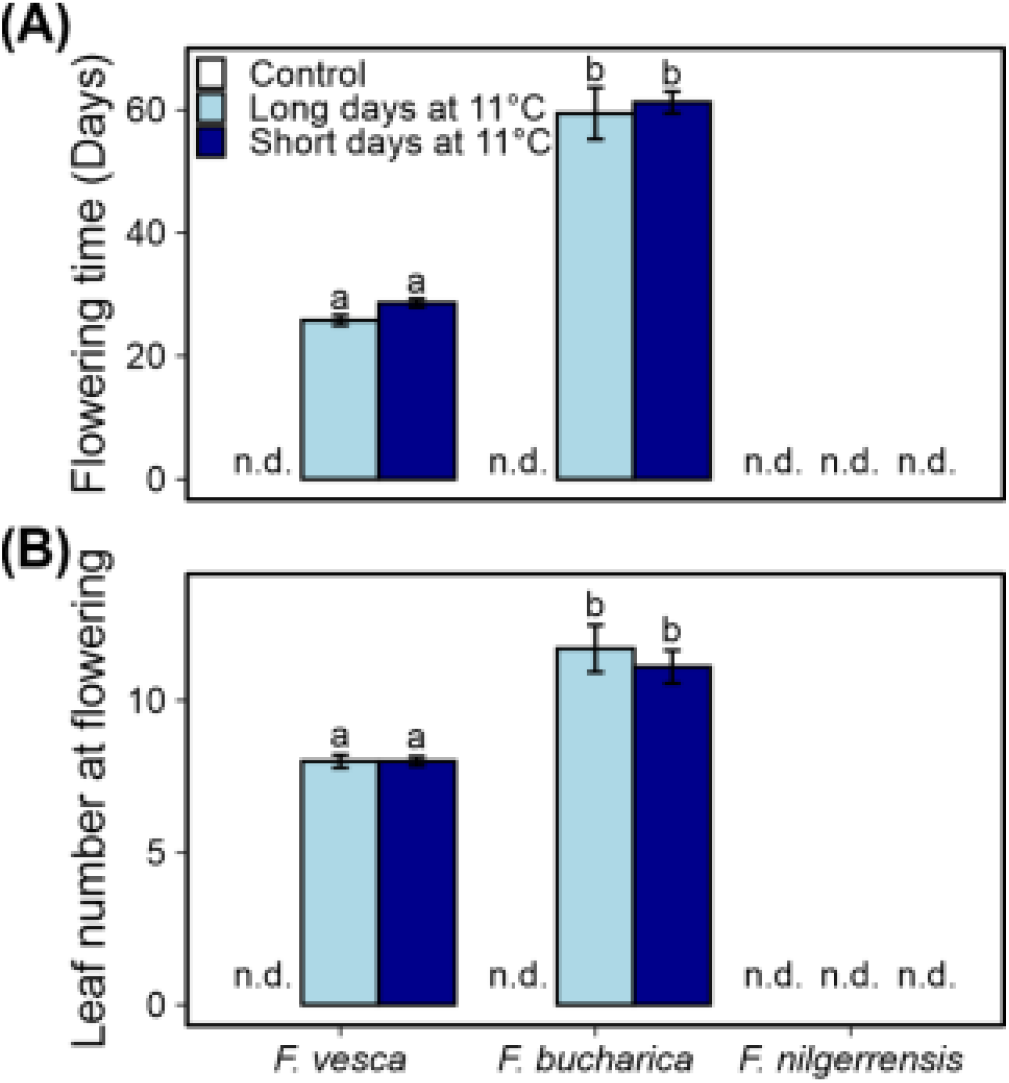
Flowering characterization of three diploid strawberry species grown under SDs or LDs at 11°C. Number of days **(A)** and number of leaves developed **(B)** until the first flower opened. Stolon-propagated plants were grown in LDs (18-h) or SDs (12-h) at 11°C for six weeks, followed by 10 more weeks in LDs (18-h) at 18°C. Control plants were grown in LDs (18-h) at 18°C from the beginning of the experiment. Flowering time was recorded every other day, starting after the treatments. Error bars represent the standard error of the mean (n=10) and different letters indicate significant differences calculated by ANOVA and Tukey’s test (P < 0.05). “n.d.” = flowering not observed during the course of the experiment (16 weeks).

Also in this experiment, we found differences in the environmental regulation on AXB fate among the three diploid strawberry species. Overall, in all species, the control plants developed stolons continuously at a relatively stable speed throughout the experiment (Figure 6A). Cool temperature strongly suppressed stolon development in *F. vesca* and stolon development did not resume after returning the plants to 18°C. Similarly to *F. vesca*, stolon development ceased also in *F. bucharica* after three weeks at 11°C under both LD and SD conditions, but in contrast to *F. vesca, F. bucharica* started stolon development again two weeks after the treatments. In *F. nilgerrensis*, stolon development was markedly slowed down at cool temperature compared with the control plants, but no clear cessation of stolon development was observed. On the other hand, *F. bucharica* and *F. nilgerrensis* plants subjected to LDs at 11°C had significantly more stolons on week nine than plants grown in SDs. In *F. vesca*, such a photoperiodic effect was not observed.

**Figure 6.**
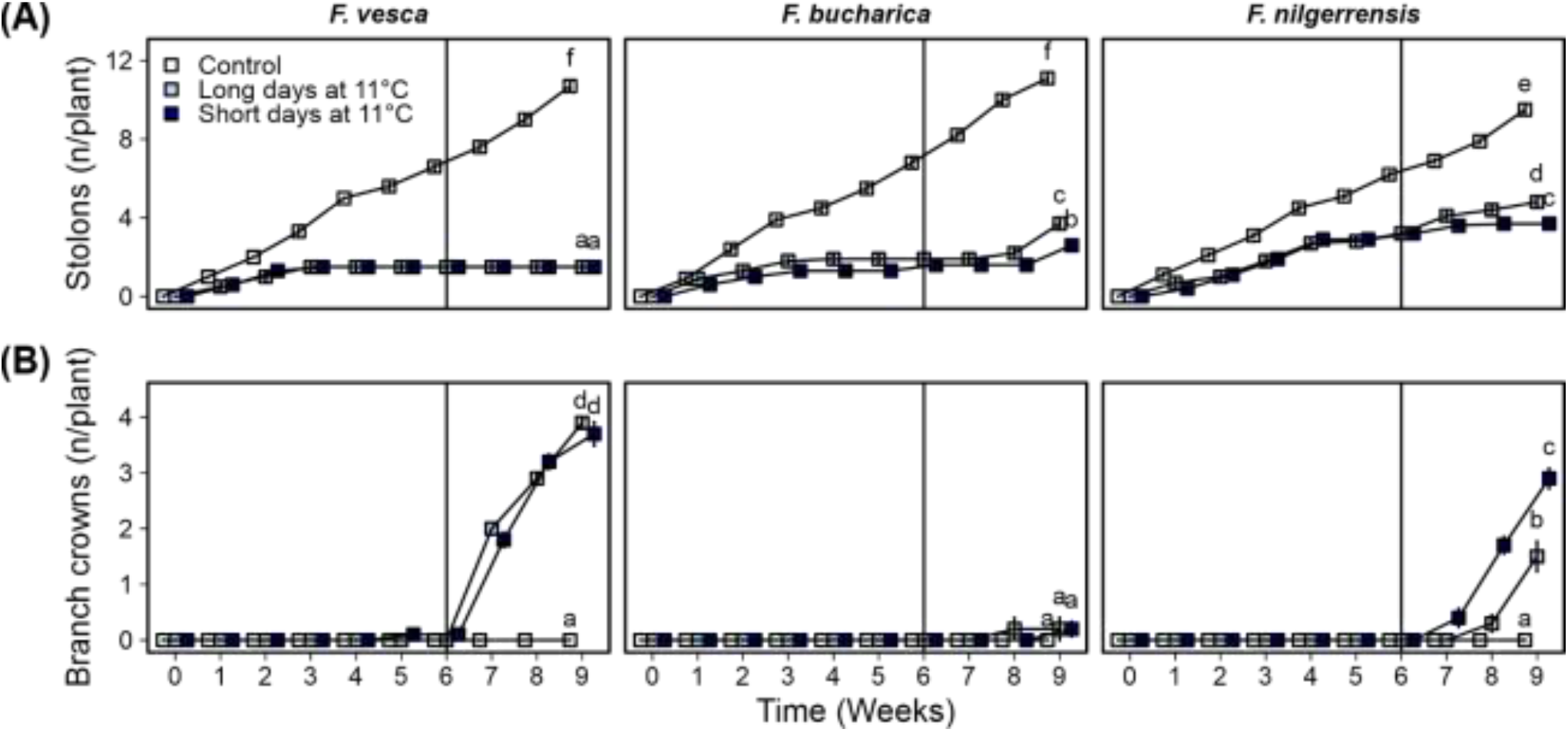
Axillary bud fate in three diploid strawberry species grown under LDs or SDs at 11°C. Number of stolons **(A)** and branch crowns **(B)** per plant. Stolon-propagated plants were grown under LDs (18-h) or SDs (12-h) at 11°C for six weeks and subsequently in LDs (18-h) at 18°C for three more weeks. Control plants were grown continuously under LDs (18-h) at 18°C. Number of stolons and branch crowns were recorded weekly. Error bars represent the standard error of the mean (n=10) and different letters indicate significant differences calculated by ANOVA and Tukey’s test (P < 0.05).

The three *Fragaria* species differed in terms of branch crown development. *F. vesca* started to develop BCs independently of the photoperiod immediately after the six-week period at 11°C (Figure 6B). On the contrary, *F. bucharica* developed very few BCs after the photoperiodic treatments at 11°C, although it behaved similarly to *F. vesca* in terms of stolon development and all the plants flowered. Also *F. nilgerrensis* started to develop BCs after the photoperiodic treatments at 11°C although plants did not flower, and this occurred about one week later in the LD treatment. At week nine, there were significantly more BCs in the SD-grown plants of *F. nilgerrensis* (Figure 6B). The control plants grown continuously under LDs at 18°C did not develop BCs in these species. Taken together, our data indicates that AXB fate in the three studied *Fragaria* species is controlled by different mechanisms; in *F. vesca*, cool temperature of 11°C promotes BC development and suppresses stolon development independently of photoperiod, while AXB fate in *F. nilgerrensis* is clearly dependent on photoperiod. Moreover, BC development in *F. bucharica* appears to be endogenously regulated, as neither photoperiod nor temperature affected BC development.

### 1.3.5 The expression of key genes correlates with flowering and AXB fate at 11°C

Next, we wanted to examine whether the phenotypical differences observed between the three *Fragaria* species at 11°C could be explained by altered expression of key genes. Earlier studies in *F. vesca* suggest that *FvSOC1* is activated by LDs at 10°C, albeit to a lesser extent than at higher temperatures (Rantanen et al., 2015). In our current experiment at 11°C, we saw clear SD-dependent downregulation of *SOC1* in *F. vesca*, as well as in *F. bucharica* (Figure 7A). In *F. nilgerrensis*, the pattern of *SOC1* expression was not very clear, although by week six the level of *SOC1* mRNA was lower in SDs than in LDs.

**Figure 7.**
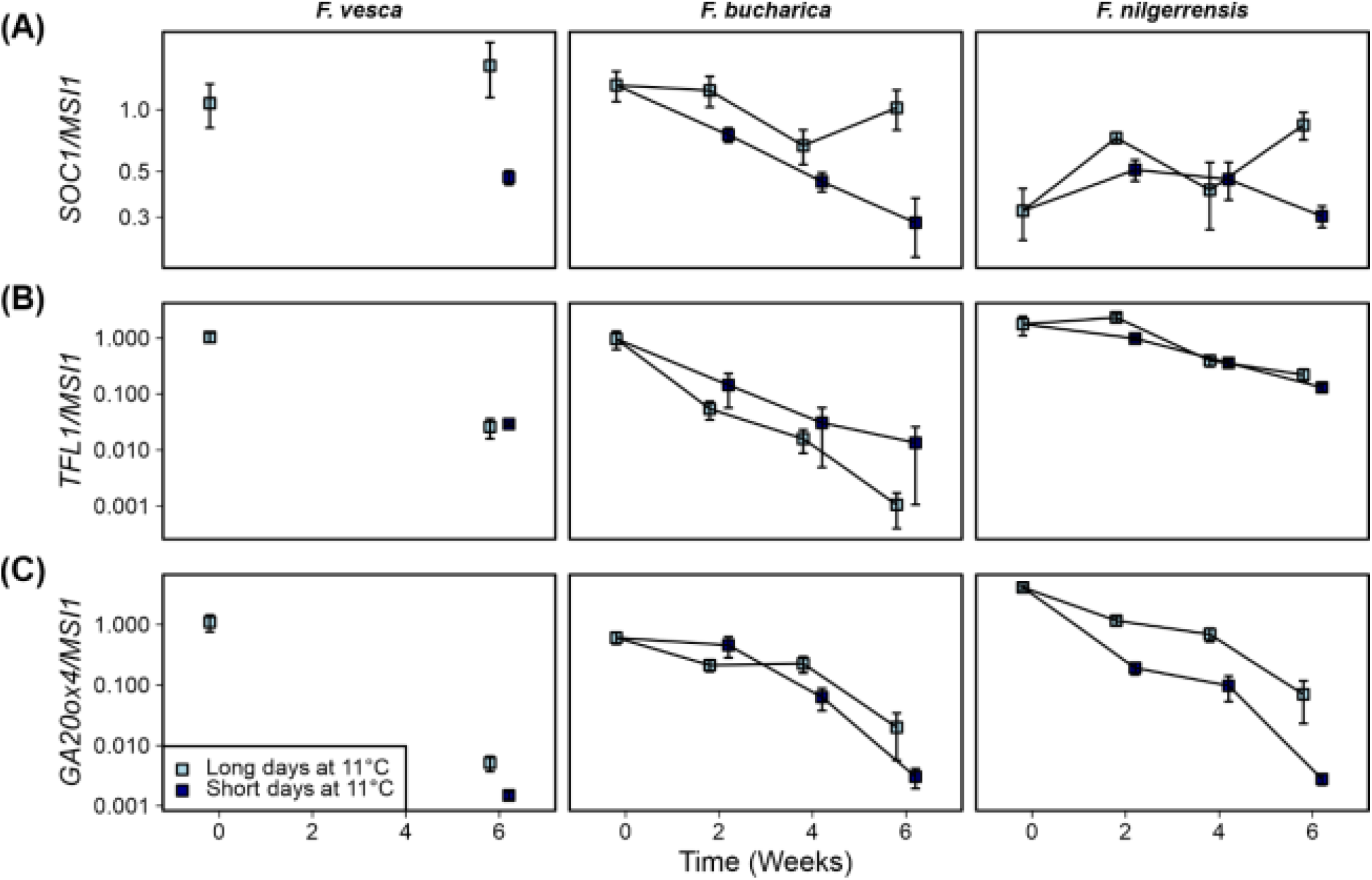
Gene expression patterns in three diploid strawberry species grown in SDs or LDs at 11°C. *SOC1* expression **(A)**; *TFL1* expression **(B)**; *GA20ox4* expression **(C)** in shoot apical samples. Stolon-propagated plants were grown in LDs (18-h) or SDs (12-h) at 11°C for six weeks. Shoot apical samples were collected at the beginning of the treatments, and 2, 4 and 6 weeks later. Week 0 *F. vesca* samples were used as calibrator for relative expression analysis. Error bars represent the standard error of the mean (n = 3–4).

As shown earlier by Rantanen et al. (2015), *FvTFL1* in *F. vesca* is de-activated independently of photoperiod at 11°C. *TFL1* was downregulated at 11°C also in our current experiment in *F. vesca*, and the expression pattern of *TFL1* was very similar in *F. bucharica* (Figure 7B). In *F. bucharica, TFL1* expression decreased to extremely low level after only two weeks, and it remained low until week 6 (Figure 7B). On the contrary, down-regulation of *TFL1* in *F. nilgerrensis* occurred slower than in *F. bucharica*. At week 6, *TFL1* expression was still higher in *F. nilgerrensis* than in the other species.

We also analysed the expression of *GA20ox4*. The effect of 11°C treatment on *GA20ox4* activity was very clear for all the species. In *F. vesca* and *F. bucharica*, the expression level was much lower after six weeks at 11°C than in the beginning of the experiment, correlating with the lack of stolon development in these two species (Figure 6A). Results in *F. bucharica* also showed that this downregulation occurred gradually in both photoperiods. By contrast, although *GA20ox4* expression in *F. nilgerrensis* also declined gradually, it was clearly more downregulated in SDs than in LDs from week two onwards. The difference between photoperiods was even more evident by week six, being in line with the effect of photoperiod on stolon development in this species.

## 1.4 Discussion

The molecular mechanisms regulating floral induction and AXB fate as a response to environmental conditions are starting to emerge in the diploid model species *F. vesca* (Hytönen and Kurokura 2020; Andrés et al., 2021). To gain a broader view on the diversity of regulation of flowering and AXB fate within the *Fragaria* genus, we studied these responses in a panel of wild diploid strawberry species originating from diverse geographical origins ranging from Asia to Europe. Some of these species inhabit very local habitats while some are widely spread around the Northern Hemisphere and are adapted to a wide range of environments (Liston et al., 2014). Here, we characterized the flowering habits of 14 accessions from seven wild diploid strawberry species under controlled environmental conditions. Based on their diverse flowering responses, we further selected two representative species, *F. bucharica* and *F. nilgerrensis*, and analyzed their flowering habits, AXB fate and expression of key genes related to these biological processes in comparison to the reference model species *F. vesca*.

### 1.4.1 Diversity of flowering responses in diploid *Fragaria* species

We discovered a diversity of flowering responses in our collection of diploid *Fragaria* species. *F. bucharica* and *F. viridis* were clearly photoperiod-insensitive at cool temperature and flowered rapidly after both SD and LD treatments at 11°C, similarly to the model species *F. vesca* (Figure 1; Rantanen et al., 2015). These results were in line with the earlier field observations on facile floral induction in these species; *F. bucharica* flowered twice during the growing season when grown in Germany (Staudt, 2006), and *F. viridis* was described as “remontant” under field conditions in South East England (Sargent et al., 2004), as also observed in the Professor Staudt Collection in Germany (data not shown). In contrast to these species, *F. iinumae, F. nilgerrensis* and *F. nubicola* #1 featured photoperiod sensitivity at 11°C; the promoting effect of SDs was obvious, with some accessions showing an obligatory requirement for SDs at even such a cool temperature. Sensitivity to photoperiod at cool temperature has been previously described in some accessions of *F. vesca*, but the effect of photoperiod was milder than in the above-mentioned species (Heide and Sønsteby, 2007). Moreover, we found LD promotion of flowering in *F. chinensis* at 11°C, a response that has not been previously described in *Fragaria* (Figure 1). Finally, *F. pentaphylla* and *F.nubicola* #2 did not flower at all after 11°C treatments. The diverse responses observed in our collection warrant further investigation to uncover mechanisms controlling flowering time variation in these strawberries.

In general, diploid strawberries flower in their original habitats in spring/summer after overwintering (Staudt, 2003; Staudt, 2005; Staudt, 2006; Staudt 2009), as well as when tested in field conditions *ex situ*. Sargent et al. (2004) characterized accessions of *F. iinumae, F. nilgerrensis, F. nipponica, F. nubicola, F. pentaphylla, F. viridis* and *F. vesca* for their flowering habits in field conditions in South East England and was able to produce flowers and fruits in all the included accessions. Likewise, a long-term exposure to 5-6°C in a greenhouse without supplemental light promoted flowering in the majority of the accessions we tested, excluding *F. iinumae* and *F. nubicola* #2 (Supplementary Table 3). In contrast, Bors and Sullivan (2005) reported that a two-month period at constant −1°C could not induce some accessions of *F. nilgerrensis, F. nubicola, F. pentaphylla* and *F. viridis* to flower and an additional floral induction treatment under 10-hour SDs at 18/15°C (day/night) was necessary. As the experimental conditions, as well as the plant materials, differ between the studies (Sargent et al., 2004; Bors and Sullivan, 2005), direct comparisons are difficult. The contrasting results highlight the need for experimentation under controlled climate to uncover the exact environmental conditions required for floral induction in *Fragaria* species. Furthermore, based on observations in the Professor Staudt Collection indicate that plant age should also be considered (data not shown).

We observed conflicting results in *F. iinumae* that was induced to flower in SDs at 11°C, but not after a long cold treatment at 5-6°C. In field experiments in Germany, as well as in its native environment in Japan and Sahalin, *F. iinumae* has been observed to flower extremely early in spring, sometimes even at sub-zero temperature (Staudt, 2005). This suggests that *F. iinumae* tolerates low temperatures extremely well, and may possess an exceptionally low temperature threshold for breaking of dormancy. In cultivated strawberry, dormancy is attained when the plants are subjected to SDs at 15°C, whereas SDs at 6°C do not induce dormancy (Sønsteby and Heide, 2006). It is possible that the cooler temperatures of 5-6°C induced dormancy in *F. iinumae*, which is why the plants did not flower during subsequent forcing in LDs (Supplementary Table 3). Further studies in *F. iinumae* are needed to uncover the specific environmental conditions required for flowering in this species.

### 1.4.2 Variation in *TFL1* regulation correlates with contrasting flowering habits in three diploid *Fragaria* species

TFL1 is a strong floral repressor in different Rosaceous species (Kotoda et al., 2006; Flachowsky et al., 2012; Freiman et al., 2012; Iwata et al., 2012; Koskela et al., 2012; Koskela et al., 2016; Charrier et al., 2019). We found a clear association between *FvTFL1* de-activation and floral induction in *F. vesca*, corroborating the earlier findings by Rantanen et al. (2015), and a similar association was found in *F. bucharica*. In both species, SDs suppressed *TFL1* and promoted floral induction at 18°C, whereas *TFL1* downregulation and floral induction occurred independently of photoperiod at 11°C (Figures 2, 4B, 5 and 7B). However, we were unable to induce flowering in *F. nilgerrensis* in these experiments, and this lack of floral induction was associated with an overall higher level of *TFL1* expression. These data suggest that, while the environmental conditions required for floral induction differ between the species, the role of *TFL1* as a floral repressor appears to be conserved in *F. vesca, F. bucharica* and *F. nilgerrensis*.

Earlier experiments in *F. vesca* identified *FvSOC1* as a major LD-activated promoter of *FvTFL1* expression at 18°C (Mouhu et al., 2013). We observed LD-dependent activation of *SOC1* expression at 18°C for *F. vesca, F. bucharica* and *F. nilgerrensis* (Figure 4A), suggesting that the photoperiodic pathway upstream of *SOC1* functions similarly in the three species. However, in *F. nilgerrensis* the pattern of *FnTFL1* expression did not coincide with that of *FnSOC1*, as *FnTFL1* activity remained at a relatively high level under both photoperiods although *FnSOC1* was strongly downregulated by SDs. This indicates an uncoupling of *FnSOC1* and *FnTFL1* expression patterns that is reminiscent of the events taking place in *F. vesca* at 23°C; at this temperature, *FvTFL1* is upregulated by an unknown pathway independently of *FvSOC1* (Rantanen et al., 2015). It is possible that the temperature threshold for the activation of this unidentified pathway is lower in *F. nilgerrensis* than in *F. vesca*, leading to photoperiod- and *FnSOC1*-independent upregulation of *FnTFL1* already at 18°C.

At cool temperature of 10-11°C, the regulatory link between *FvSOC1* and *FvTFL1* is abolished in *F. vesca;* at this temperature, *FvTFL1* is de-activated independently of photoperiod even though *FvSOC1* expression is still photoperiodically regulated (Rantanen et al., 2015). We observed the *SOC1*-independent downregulation of *TFL1* in our current experiment conducted at 11°C for both *F. vesca* and *F. bucharica*, while *F. nilgerrensis* differed from the other two species by showing much slower *TFL1* downregulation (Figure 7B). Approximately 60% of *F. nilgerrensis* plants flowered after 6-week treatment in SDs at 11°C in the initial screening (Figure 1) and after being subjected to a long cold treatment at 5-6°C (Supplemetary Table 3). This suggests that *F. nilgerrensis* may require a longer treatment at 11°C or photoperiods shorter than 12 hours to gradually downregulate *FnTFL1* to the level that permits flower induction to occur. In consistence with this hypothesis, the species originates from low latitude area with relatively mild seasonal changes of temperature and photoperiod, where the period of flower inductive conditions is longer than in the colder habitats of *F. vesca* and *F. bucharica* (Liston et al., 2014).

In conclusion, *TFL1* is a key integrator of environmental signals in the three studied diploid strawberry species, and variation in the downregulation of *TFL1* may explain the observed differences in their photoperiodic and temperature responses. Further studies are needed to explore what are the molecular mechanisms controlling variation in *TFL1* regulation.

### 1.4.3 GA20ox4 promotes stolon development in the three *Fragaria* species

Although the regulation of AXB fate in *F. vesca* has received attention in the recent years (Tenreira et al., 2017; Caruana et al., 2018; Li et al., 2018; Qiu et al., 2019; Andrés et al., 2021), AXB fate regulation has remained unexplored in other wild *Fragaria* species. We found that, in line with previous studies in *F. vesca* (Mouhu et al., 2013), SDs at 18°C completely inhibited stolon development in *F. vesca* and *F. nilgerrensis* after three and six weeks of treatments, respectively. However, stolon development of *F. bucharica* was reduced only slightly under SDs (Figure 3B). In *F. vesca*, this photoperiodic response is mediated via *FvSOC1* that promotes stolon formation in LDs by upregulating *FvGA20ox4* in AXBs, and the downregulation of these genes stops stolon formation (Andrés et al., 2021). In our current experiment, *GA20ox4* expression was in line with that of *SOC1* in *F. vesca* and *F. nilgerrensis* at 18°C, but not in *F. bucharica* that exhibited clear photoperiodic regulation of *SOC1* but barely noticeable differences in *GA20ox4* expression (Figure 4). Andrés et al. (2021) found that, at 23°C, *FvGA20ox4* expression is promoted by an unknown factor in SD-grown *F. vesca* plants, in spite of downregulation of *FvSOC1*. Perhaps the same factor upregulated *GA20ox4* at 18°C in SD-grown *F. bucharica* plants in the current experiment.

Corroborating with earlier studies in *F. vesca*, stolon development ceased in all three species at 11°C regardless of the photoperiod, but this happened later in *F. nilgerrensis* than in other species (Figure 6A; Heide and Sønsteby, 2007; Andrés et al., 2021). The cessation of stolon development was associated with gradual downregulation of *GA20ox4* expression in all three species indicating that *GA20ox4* controls stolon development also in *F. bucharica* and *F. nilgerrensis* in these conditions. This finding is in line with the result of Andrés et al. (2021), who observed photoperiod-independent downregulation of *FvGA20ox4* in the *F. vesca* accession ‘Hawaii-4’ at cool temperature. It is notable that the expression of *GA20ox4* does not follow the expression pattern of *SOC1* at 11°C, not in our current experiment (Figure 7), nor in the earlier study by Andrés et al. (2021). These data indicate that, at 11°C, *GA20ox4* expression is regulated by factors other than *SOC1* in the three species.

### 1.4.4 Different pathways regulate branch crown development in the three species

Although flower-inductive treatments promoted BC development in *F. vesca*, BC formation did not correlate with flowering in *F. bucharica* and *F. nilgerrensis*, highlighting the diverged regulation of BC development in the three species. In *F. nilgerrensis*, SDs and cool temperature activated BC development independently of flowering, which was previously found also in late- and non-flowering *F. vesca* mutants with high *FvTFL1* expression levels (Andrés et al., 2021). In *F. bucharica*, none of our tested conditions could promote BC development, although we witnessed clear photoperiodic and temperature regulation of flowering (Figure 3B and 6B). In *F. bucharica* grown at 18°C, the lack of BCs was consistent with high *FbGA20ox4* expression level in both SDs and LDs, and only flowering forced the topmost AXB to continue the growth of the leaf rosette sympodially. At 11°C, however, BCs were missing regardless of the downregulation of *FbGA20ox4* and the cessation of stolon formation. These findings suggest that BC development in *F. bucharica* was inhibited by factor(s) other than apical dominance, even after the downregulation of *FbGA20ox4*. This contrasts with findings in a stolonless *ga20ox4* mutant of *F. vesca* that exhibits strong apical dominance, and forms BCs only after flower induction or decapitation of the shoot tip (Andrés et al., 2021). Also columnar apple trees show remarkably strong apical dominance, and almost all of their axillary shoots re reproductive spur shoots (Kelsey and Brown, 1992), analogous to *Fragaria* BCs. These trees have ectopic *MdDOX-Co* expression in shoots, in addition to its normal root-specific expression pattern, which hampers GA biosynthesis and leads to columnar phenotype (Okada et al., 2020; Watanabe et al., 2021).

Striking differences in BC development of *F. bucharica* and *F. nilgerrensis* may be adaptations to their native habitats. *F. bucharica* is found at high altitude regions with very short growing season in Himalayas (Hummer et al., 2011; Johnson et al., 2014). Such alpine areas are usually dominated by species with low reproductive vigor that favor vegetative propagation over the comparatively riskier sexual reproduction (Billings 1968; Eriksson 1997; Klimes et al., 1997; Grime 2002; Zhang et al., 2010). Therefore, the lack of branch crowns that limits the number of inflorescences to maximum one per plant may be an adaptive trait that makes vegetative reproduction through stolons a primary reproductive mode in *F. bucharica. F. nilgerrensis*, in contrast, is native to habitats with very mild winters and long growing seasons in the Southeast Asia. Such conditions are more suitable for sexual reproduction, which may explain why this species develops abundant BCs in SDs in autumn/winter to enable the plant to form several inflorescences during the following growing season.

## 1.5 Conclusions

In this work, we provide the first phenotype and gene expression level analyses on the control of flowering and axillary meristem fates in several wild diploid *Fragaria* species under controlled environmental conditions. We show that the examined species feature a wide range of flowering responses, and changes in *TFL1* regulation is the key to understanding the different responses. Moreover, we show that the environmental regulation of *GA20ox4* varies in the three studied species, and this variation is associated with differences in stolon development. Finally, our results on BC development suggest diverged regulation of this process in *F. vesca, F. bucharica* and *F. nilgerrensis*. To summarize, the diverse phenotypical responses provide an excellent starting point for carrying out further experiments to elucidate the genetic bases of these responses. We anticipate that such studies would provide new means to control yield formation in Rosaceous fruit and berry crops through 1) altered flowering characteristics based on modifications of *TFL1* regulation, and 2) improved plant architectures by optimizing the balance between the formation of short and long shoots.

## 3 Author Contributions

GF, JA, EK and TH designed the experiments. KO provided diploid *Fragaria* species plant materials and participated in the selection of genotypes for the study. GF and JA carried out the experimental work, statistical analyses and drafted the manuscript. EK and TH supervised the work and edited the manuscript. All authors commented on and accepted the final manuscript version.

## 5 Acknowledgments

Doctoral Programme in Plant Sciences at the University of Helsinki is acknowledged for JA’s salaried PhD student position. We are also grateful to MSc student Kaiyue Qin about her help with phenotypic observations.

